# High-Resolution 3D Histology of the Murine Kidney Using Synchrotron X-Ray Micro-CT

**DOI:** 10.64898/2026.02.11.705367

**Authors:** Humberto C. Joca, Pablo W. Aguiar e Silva, João V. Ribeiro dos Santos, Elayne V. Dias, Thais P. Barbosa, Karina Y. Degaki, Raphael M. Neto, Maiara F. Terra, Renata S. Rabelo, Mateus B. Cardoso, Ângela Saito, Thayná M. Avelino

## Abstract

Conventional two-dimensional (2D) histology relies upon destructive sample preparation and stereological estimation, frequently leading to sampling bias and loss of critical spatial context required for understanding renal structure relationships. Here, we detail a novel pipeline for high-resolution 3D histology of *ex vivo* murine kidneys using X-ray micro-computed tomography (micro-CT) at the high flux of a synchrotron light source, the architecture of the nephron and associated microvasculature necessitates three-dimensional (3D) analysis to accurately characterize its complexity. Soft-tissue contrast was optimized through an established phosphotungstic acid (PTA) staining protocol, enabling robust mapping of macro and microstructures via absorption contrast. Multi-scale imaging was performed, providing whole-organ context at resolutions around 3 μm and achieving sub-micron detail (down to 400 nm) in targeted regions of interest (ROI) of the renal cortex. Utilizing machine learning segmentation pipelines optimized for large volumetric datasets, we extracted crucial 3D quantitative morphometric data. The results presented herein demonstrate accuracy and morphological insight achievable through synchrotron-based 3D imaging, establishing a robust method for quantitative preclinical research.

## 1. INTRODUCTION

Historically, research on morpho-physiology relied heavily on conventional two-dimensional (2D) histology using traditional staining methods such as H&E and Masson’s Trichrome staining^1^. This approach remains invaluable for providing morphological details for research and (pre-)clinical diagnosis due to its simplicity, ease of access, cost-effectiveness, and repeatability. However, despite its undeniable usefulness and widespread adoption, this approach has a fundamental limitation: it generates only 2D images by destructively preparing a three-dimensional (3D) tissue sample. This process inevitably compromises the study of overall structural architecture and introduces sampling bias, especially when attempting to quantify parameters that require a complete volumetric context, which dictates function^3^. That demands a paradigm shift toward a quantitative 3D analysis to fully resolve the biological tissue complexity.

X-ray computed tomography (CT) offers a powerful solution for the non-destructive, 3D morphological investigation of biological samples. Specifically, micro- and nano-computed tomography (micro-CT and nano-CT) enable rapid imaging of internal and external structures at resolutions suitable for detailed morphological studies^1^. However, the application of CT to soft biological tissue is limited by the lack of contrast within the tissue due to its low X-ray attenuation coefficien^3^. To overcome this inherent limitation, a few approaches can be used, such as contrast enhancement via heavy-metal staining, phase contrast, and/or a high-flux X-ray source^4,5^. Contrast enhancement can be achieved through heavy-metal staining to highlight histological targets, such as the cytoplasm or extracellular matrix, and better visualize morphological detail^5,6^. Simultaneously, obtaining high spatial resolution (approaching 1 μm or better) would require a powerful X-ray source. Synchrotron radiation can provide the high X-ray flux and spatial coherence needed to balance high spatial resolution with an adequate signal-to-noise ratio (SNR) to allow fast acquisition^4^. This is highly desirable for high-throughput preclinical studies, as conventional laboratory benchtop micro-CTs often require extended scan times and are limited to lower effective resolutions.

Here, we use murine kidneys as biological tissue to establish and validate a rigorous 3D histology pipeline. The mammalian kidney maintains body homeostasis through a dense and functionally compartmentalized structure. Its function is intrinsically dependent on the precise, complex 3D spatial relationships among the nephrons, the microvasculature, and the elaborate tubular system^2^. Consequently, many renal diseases, particularly Chronic Kidney Disease (CKD), manifest as volumetric and spatial alterations, such as nephron loss, microvascular degradation, and tubulointerstitial fibrosis^7,8^. Murine models of CKD, including chronic kidney injury models, are routinely employed to investigate these pathological changes and assess potential interventions and would benefit from 3D morphological investigation provided by high-resolution micro-CTs^2,7–9^.

In this study, we optimize a contrast enhancement protocol for renal soft tissue imaging, specifically demonstrating its ability to enhance cytoplasmic and interstitial structures for detailed 3D visualization and demonstrate the multi-scale imaging capability of the synchrotron X-ray tomography, enabling efficient transition from whole-organ context (macro-scale, up to 3 μm) to sub-micron region of interest analysis (micro-scale, below 500 nm) utilizing the unique capabilities of the Mogno beamline at the Sirius Synchrotron Light Source to enable fast acquisition while maintaining high spatial resolution^10^. We also developed and applied a semi-automated pipeline using machine learning tools to process the resulting volumetric data and extract accurate, quantitative 3D morphometric indices of murine kidney for potential use in pre-clinical studies of renal pathophysiology.

## 2. METHODS

### 2.1. Animals

All procedures involving experimental animals adhered strictly to institutional (CEUA-CNPEM), Brazilian Council for the Control of Animal Experimentation (CONCEA) and international guidelines (NIH Guide for the Care and Use of Laboratory Animals) for the care and use of laboratory animals. The methods employed were designed to minimize pain, suffering, and discomfort, utilizing proper anesthesia and analgesia throughout any procedures and euthanasia. All experimental procedures were performed by trained individuals, and the entire study design, including housing and husbandry of the murine models (Neonatal and adult C57BL/6), was approved by Ethics Committee on the Use of Animals of the Brazilian Center for Research in Energy and Materials (CEUA-CNPEM #126/2024 and #143/2025) prior to the initiation of the study.

### 2.2. Sample Preparation and Contrast Enhancement

Following euthanasia, organs (kidney, lungs, heart) from neonatal C57BL/6 mice (day 0) were removed and fixed with 4% paraformaldehyde (PFA) by immersion (24h – 4°C). For adult C57BL/6 mice (10-12 weeks-old), the whole animal was perfused *in situ* by a cannula inserted into the left ventricle. Using a syringe (20ml ∼ 1ml/min), heparinized PBS solution was perfused followed by 4% PFA in PBS solution for fixation. Following the fixative perfusion, the kidneys were then harvested and post-fixed overnight in 4% PFA (4°C). After fixation, murine organs were washed using PBS (3 times) before being further processed.

The usage of contrasts was critical. As soft tissues inherently lack sufficient X-ray absorption for effective imaging, we tested different contrast agents to select the appropriate contrast enhancement^6^. We used iodine-based Lugol’s solution, and Phosphotungstic Acid (PTA, 1%) to contrast neonatal kidneys, heart and lungs overnight. After the staining, the samples were dehydrated stepwise, from 50% to 100% ethanol. For the whole organ, prior to X-ray tomography, the samples were mounted inside pipette tips (1000ul) filled with 100% ethanol then sealed with epoxy, to prevent leakage and evaporation, and kept at 4°C.

For the adult kidneys, longitudinal sections of kidney (1 mm) were cut using a vibratome (Leica VT1000S) and samples were extracted from renal cortex using a punch (1 mm diameter). The samples (1 x 1 mm cylinders) were dehydrated to 70% ethanol, contrasted with PTA (1% in 70% ethanol, overnight) then further dehydrated stepwise to 100% ethanol. Since these samples are for sub-micron region of interest analysis, they were embedded in paraffin prior to X-ray zoom tomography for better mechanical stability during imaging. After the dehydration, the samples were immersed in an ethanol:xylene (1:1) solution, followed by two 15-min incubations in xylene. Paraffin infiltration was initiated with a 20-min incubation in a xylene:paraffin (1:1) mixture and completed with two additional paraffin baths of 5 min each. The paraffin-embedded samples were allowed to solidify inside a silicone mold at 4 °C overnight, after which the solid block was removed and trimmed to approximately 2 mm width and 5 mm height and kept at 4°C until the day of x-ray imaging.

### 2.3. Synchrotron X-Ray Micro-CT

Imaging was performed *ex vivo* using the cone-beam tomography setup at the high-flux Mogno beamline of the Sirius Synchrotron Light Source (CNPEM, Campinas-SP, Brazil). Mogno beamline is equipped with experimental nano- and microtomography stations^11^, featuring an indirect detection system based on a sCMOS camera (pco.edge 4.2) and a microscope with multiple lenses (2X and 5X). The Mogno beamline optical configuration enables variable field-of-view (FOV) and spatial resolution, crucial for multi-scale biological imaging of both the overall organ structure (e.g., cortex and medulla) and histology (e.g., tubules, micro vessels, glomeruli). X-ray tomography was performed using synchrotron light source (Mogno beamline, Sirius, Brazil) with a 3.2 T superbend, beam size of 22.1 x 8.5 mm^2^ (HxV, rms), and pixel size of 3.6 μm for whole neonatal organs or 353 nm for zoom tomography using renal cortex samples (adult mice). 2,048 X-ray transmission images were acquired by rotating the sample to a full 360 degrees around a fixed axis at uniformly spaced angular steps. These images were used to generate a stack of sinograms, which were then computationally transformed into a 3D density map.

### 2.4. Histology

A subset of samples of adult kidneys embedded in paraffin was sectioned using a microtome at 5 μm sections (Leica HistoCore AUTOCUT) and stained with H&E for comparison with 3D tomography slices. Briefly, slides with paraffin-embedded sections are baked and then immersed in xylene for deparaffinization and gradually rehydrated through a series of decreasing ethanol concentrations (100 to 70%) and finally distilled water. Since the tissue was previously treated with PTA, it requires treatment with a base (0.1 M NaOH for 30 minutes) after rehydration to remove the PTA and allow proper H&E staining^12^. After the base is washed with distilled water, tissue undergoes nuclear staining using Harris Hematoxylin (Sigma), and cytoplasmic staining with alcoholic Eosin (Sigma). Following dehydration using increasing ethanol concentrations and clearing in xylene, the slides are mounted with a coverslip and mounting media (DPX, Sigma). 12 images (3×4 – 10% overlap) were acquired to capture the whole sample using a 20X objective (Leica DM6 Microscope) equipped with a color camera (Leica DFC7000 T, 1.25X camera port). The microscope and camera controls, images acquisition, and image stitching to single histological image were performed using LAS X software (version 3.6.0, Leica).

### 2.5. Full Sample Segmentation Pipeline for Quantitative Analysis

The 3D reconstructed data of the full sample, acquired from the Mogno beamline, was post-processed using 3D Slicer equipped with nnInteractive, an AI-assisted module. A segmentation pipeline was developed based on PTA staining contrast features, through which the reconstructed volumes went through a specific series of processing steps.

The identified structures were categorized into two groups: microstructures (such as glomeruli, tubules, and small vessels) and macrostructures (such as the capsule, medulla, and ureter). These structures were partially annotated manually to establish the relevant morphological features during the processing. For instance, glomeruli present high intensity with a sharp intensity drop-off at the edges, delineating the capsule area. On the other hand, tubules are characterized by regions of very low intensity (representing the lumen) bordered by membranes of medium intensity, allowing for the distinction of their boundaries amid numerous adjacent tubular structures.

Following the definition of structural features, image processing techniques were applied to enhance data quality. Initially, background and irrelevant data were removed from the main sample region using a global threshold. Afterwards, an ambient occlusion algorithm was applied to this threshold label map to highlight cavities and delineate the boundaries of the main body (Figure 3). A second thresholding step was then performed specifically on these boundaries, effectively isolating the sample and eliminating external noise. This method also aided in the attenuation of ring artifacts, as most of these artifacts were located outside the strictly defined sample boundary.

These preprocessing steps, along with filter applications, served as the baseline for the segmentation process. The first structures targeted for segmentation were the glomeruli. Initially, a median filter was applied to reduce noise and sharpen boundaries. However, due to the presence of surrounding structures with similar intensity values, standard global or local thresholding methods proved insufficient for accurate labeling. Consequently, an AI-assisted annotation approach was selected.

Given the large size of the original image, full manual annotation was not feasible. Instead, a representative sub-volume was cropped and used as a dataset to train the AI model using the nnInteractive^13^ module in 3D Slicer for glomeruli labels annotation. This tool utilizes an internal U-Net architecture that processes user input prompts, such as points and scribbles, to infer the full segmentation of a structure. While this process can be challenging in low-contrast regions, it is highly effective for large tissues and continuous structures. For repetitive structures with consistent morphological patterns, like the glomeruli, this approach significantly accelerated the annotation workflow (Figure V1). Since glomeruli exhibit low morphological variation, they constitute an ideal dataset for AI training.

For the training phase, the Biomedisa^14^ software was employed. It provides both programming and CLI interfaces, accepting annotated labels as input and allowing for the configuration of various hyperparameters, including data augmentation operations, epoch count, validation steps, and U-Net layer filters.

The model was trained using an NVIDIA A100 (40GB) GPU. Training for 40 epochs took approximately 25 minutes, while inference on the full image volume was completed in just 4 minutes. Despite the use of a limited dataset, derived from a crop of the target image itself, the inference output performance was remarkable. Only minor morphological operations, such as closing, erosion, and the removal of small noise islands, were required to ensure the segmentation boundaries were smooth and topologically consistent (Figure V.1).

### 2.6. Statistics

All the data are presented as mean value + standard deviation (SD). All descriptive statistical analyses and data visualizations were performed using R version 4.5.1. Data wrangling, visualization summary statistics were performed using the tidy verse suite, including dplyr and ggplot2 packages.

## 3. RESULTS

### 3.1. Phosphotungstic Acid for Contrast Enhancement of Soft Tissues

Lugol- or PTA-enhanced micro-CT imaging successfully provided contrast for visualization of the macroscopic architecture of neonatal murine organs (Figure 1). Overall, PTA staining yielded superior image quality compared to Lugol’s solution, providing higher contrast and improved delineation of anatomical features. PTA-enhanced samples allowed detailed visualization of the whole heart (Figure 1D), lung (Figure 1E), and kidney (Figure 1F), whereas the corresponding Lugol-contrasted organs (Figure 1A–C) exhibited lower contrast and reduced structural definition. Among the analyzed organs, the kidney sample showed the most favorable outcome, with cellular-level details and high-fidelity volume renderings (Figure 2). This highlights the efficacy of PTA as a contrast agent in microtomography imaging and points to the kidney as a promising model for more in-depth image data processing and segmentation.

**Figure 1.**
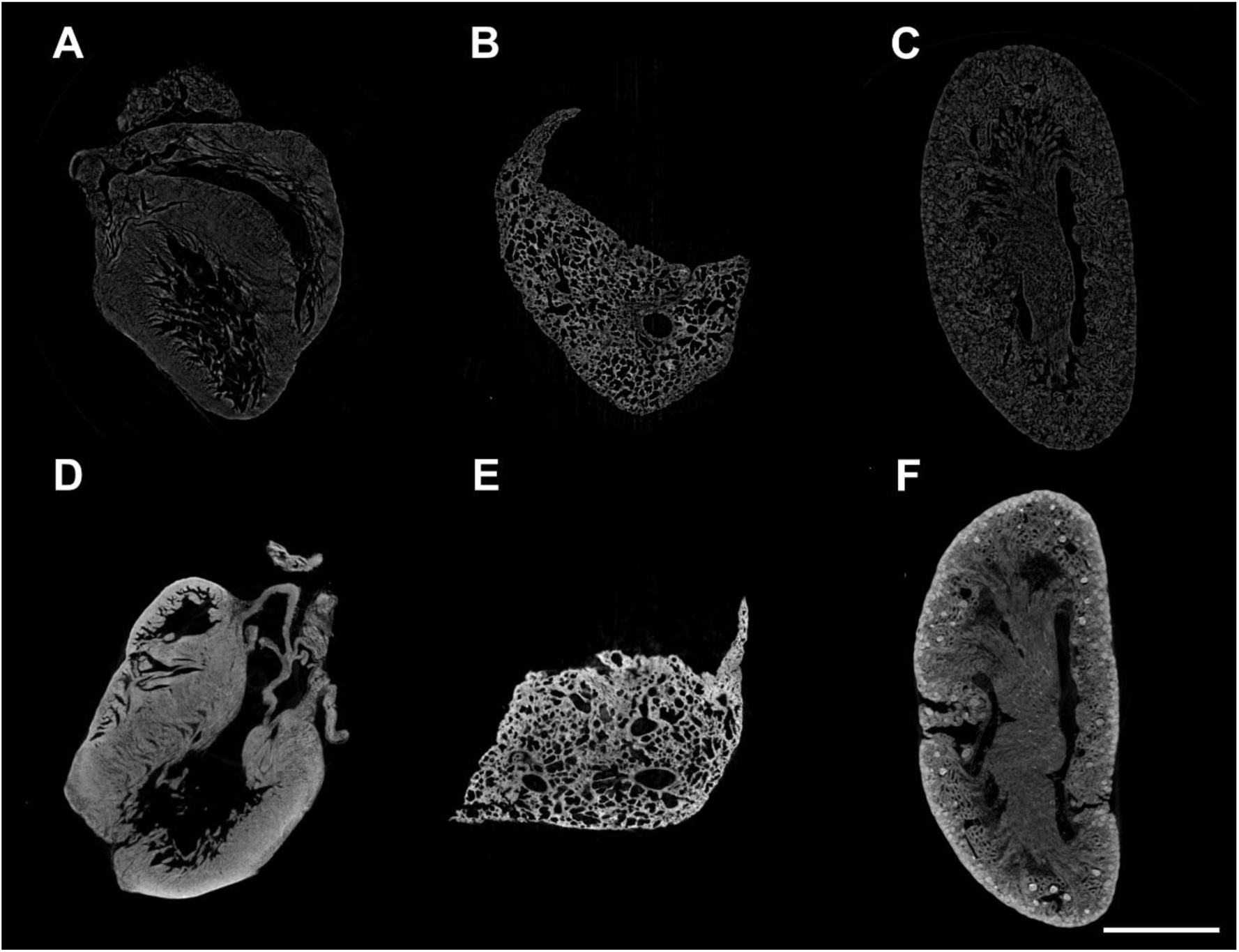
Contrast enhancement of soft tissues from neonatal murine heart (**A**,**D**), lungs (**B**,**E**), and kidney (**C**,**F**). Phosphotungstic Acid (PTA) stained samples (**D**,**E**,**F**) shown higher contrast when compated to Lugol’s solution-stained samples (**A**,**B**,**C**), allowing detailed visualization of the organ’s structure. Calibration bar: 1mm.

**Figure 2.**
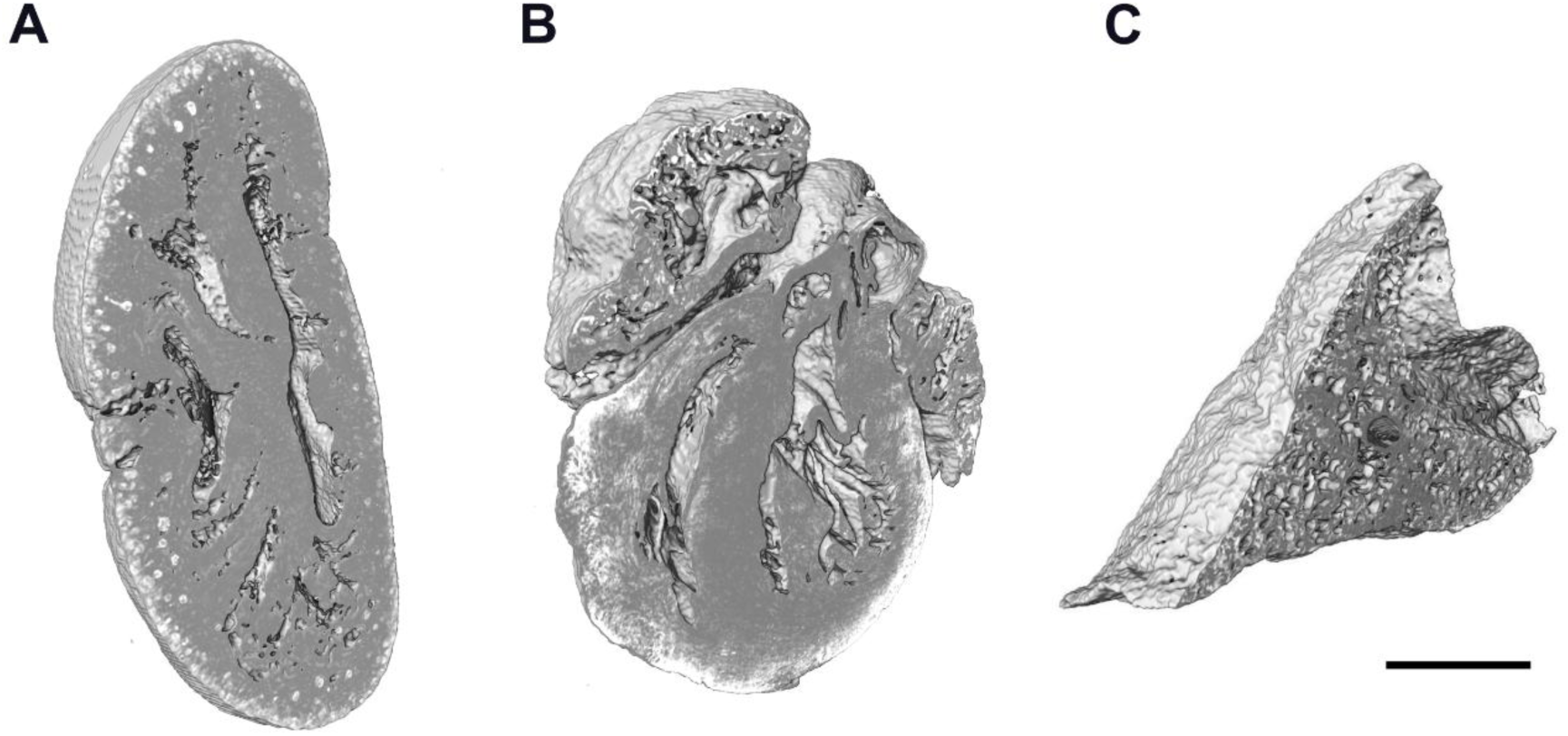
Phosphotungstic Acid (PTA) stained organs allowed high-fidelity volume renderings from neonatal murine kidney (**A**), heart (**B**), and lungs (**C**) with details at a cellular level. Calibration bar: 1mm.

### 3.2. Renal macro-structure resolved by synchrotron micro-CT

Whole kidney microtomography datasets were subjected to a multistep preprocessing workflow to reduce imaging artifacts, suppress background signal, and minimize noise while preserving fine structural details (Figure 3; see Section 2.5 for details). Raw volumes were first corrected for acquisition-related artifacts and non-uniform background intensity, followed by noise reduction to improve overall signal-to-noise ratio (Figure 3A–C).

Due to the strong and homogeneous X-ray absorption provided by PTA staining, densely packed tubular structures and the surrounding renal parenchyma exhibited high intrinsic contrast (Figure 3A). This contrast enabled effective enhancement of specific anatomical features by applying structure-preserving filters, including median filtering for noise suppression and both the black-top-hat and Frangi vesselness filters to emphasize tubular and lumen-containing structures (Figure 3B).

**Figure 3.**
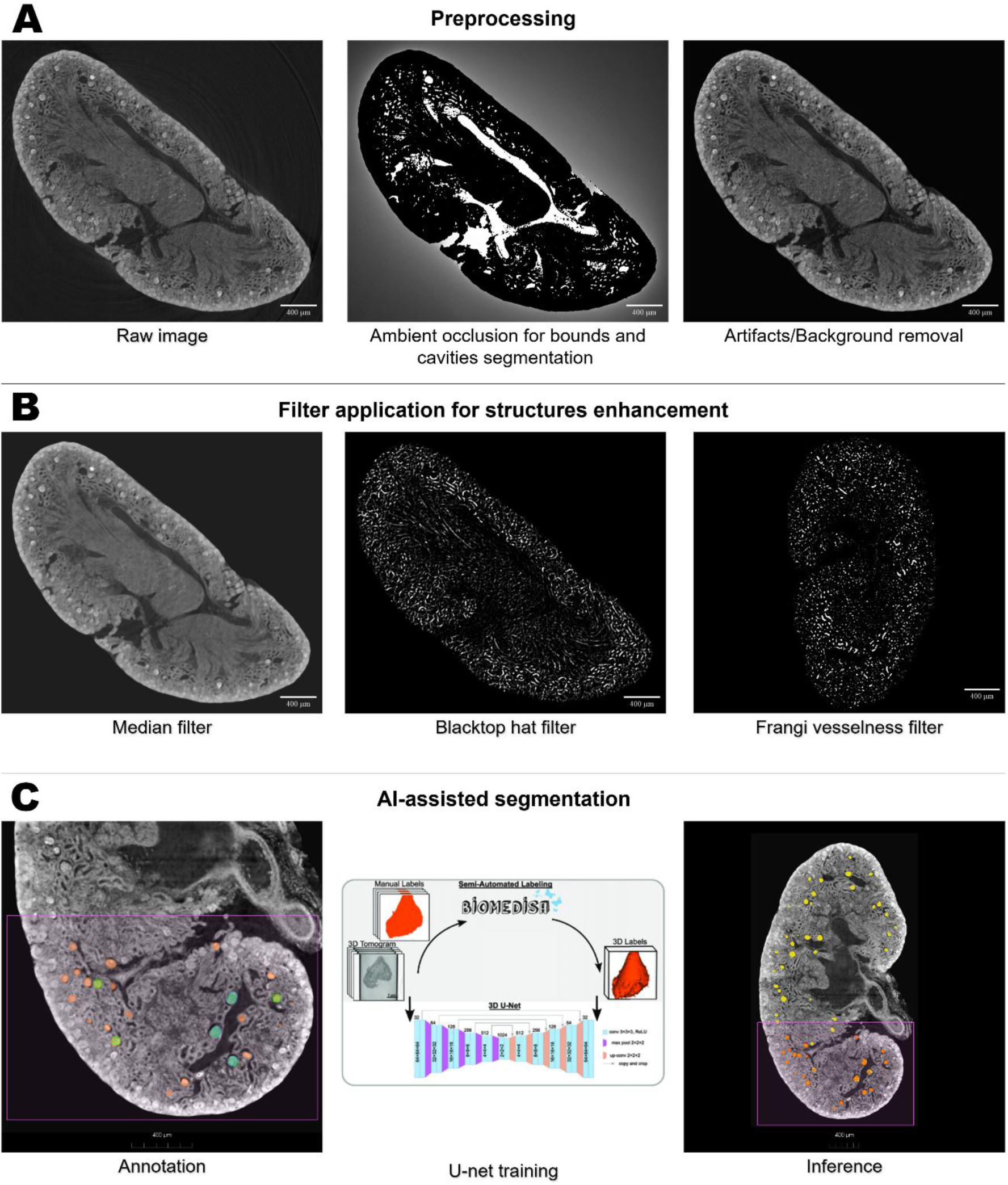
Segmentation pipeline for quantitative analysis involves 3 major steps, pre-processing for artifacts and background removal, application of filter to enhance morphological features for easier segmentation, and AI-assisted segmentation using interactive annotation with machine learn prediction of the structures to be segmented (see Methos for details).

The resulting preprocessed volumes were well suited for semi-automated interactive segmentation (Figure 3C) using nnInteractive, an open-set, 3D deep-learning–based segmentation framework trained on diverse volumetric datasets and smart interpolation of segmented mask using Biomedisa, diffusion algorithm developed for synchrotron X-ray microtomography datasets^14,15^.

This approach allowed efficient delineation of renal compartments with minimal manual annotation, providing a robust foundation for downstream three-dimensional reconstruction and quantitative analysis. Volume renderings (Figure 4) derived from those global context scans (3.6 μm pixel size) and segmented areas clearly delineated the renal capsule (Figure 4Ai), cortex (Figure 4Aii), and medulla (Figure 4B), providing an essential geometric framework for orientation and localization of the renal structures. Furthermore, this approach was able to successfully segment the glomeruli and tubules (Figure 4C). More specifically, we obtained volumetric measurements for glomeruli (Figure 4D, total count: 1015; volume: 101.13 ± 64.46 x10^3^ µm^3^, diameter: 55.65 ± 10.9 µm) and for renal cortex (4,9 mm^3^ - glomerular density: 206 glomeruli / mm^3^) and medulla (1,8 mm^3^ - cortex-to-medulla ratio: 2.7). Those macromorphological parameters can be used for histopathological evaluation in pre-clinical studies.

**Figure 4.**
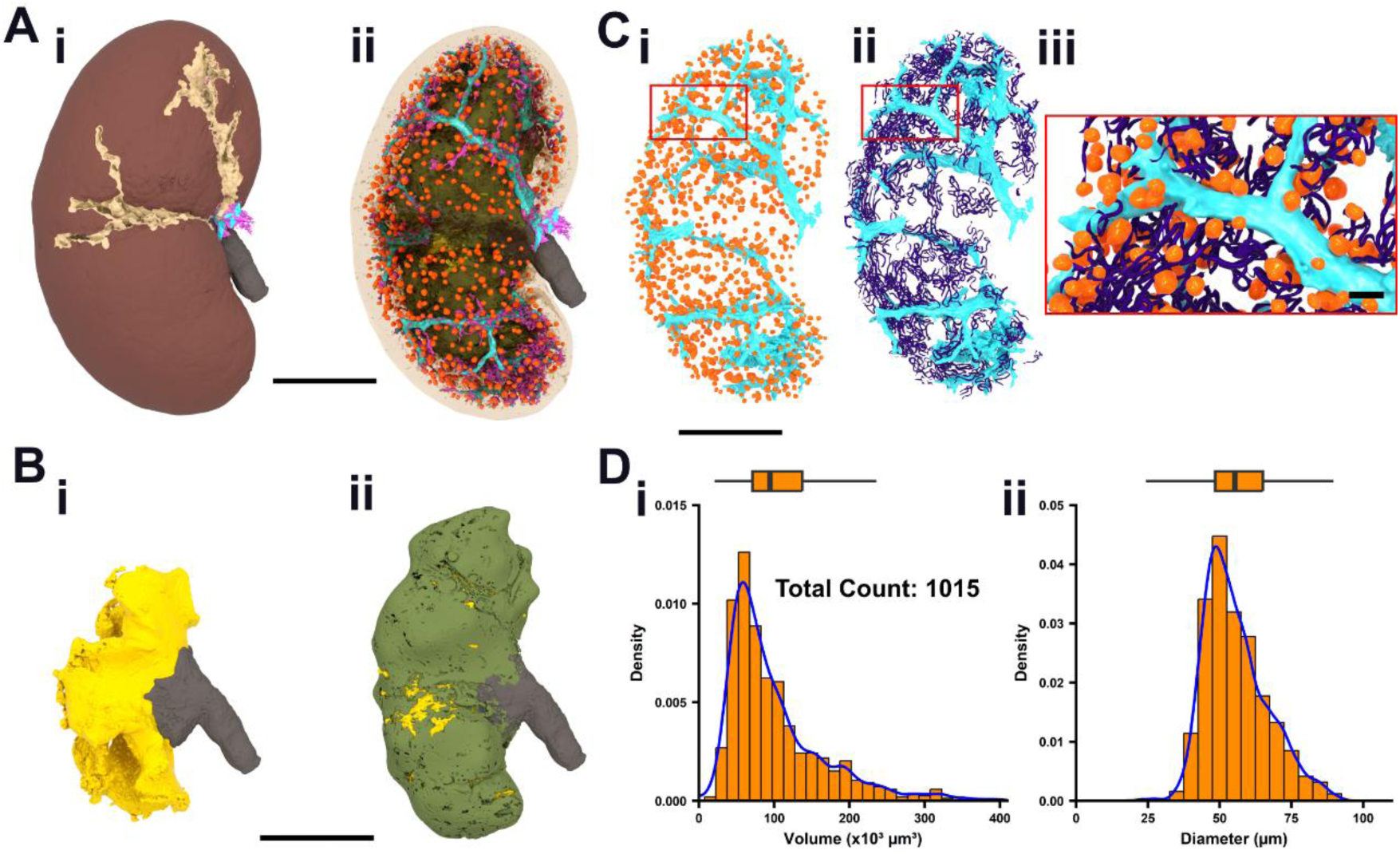
Volume renderings from whole neonatal murine kidney. Segmented areas clearly delineate the renal capsule (dark brown, **Ai**), cortex (light brown, **Aii**), pelvis (yellow, **Bi**), medulla (green, **Bii**), ureter (dark gray, **B**) and large vessels (light blue, **C**). The pipeline also successfully segmentation of smaller structures like glomerulus (orange; **Ci**), tubules (purple; **Cii**) both (**Ciii** – Zoomed area of Panel C renderings) and obtain volumetric measurements for each individual glomerulus within the cortex (**D**). Calibration bar: 1mm (**A**,**B**,**Ci** and **Cii**); 500 µm (**Ciii** inset).

### 3.3. Renal sub-micron details by zoom tomography

The multi-scale approach proved critical for correlating organ-level pathology with cellular-level detail. Targeted high-resolution ROI scans (below 500 nm pixel size, Figure 5A) demonstrated the ability to resolve fine morphological characteristics necessary for histopathological diagnosis. This ability to non-destructively visualize cellular architecture in 3D establishes the technique as a true volumetric histology approach. Using similar approach to whole organ scans, the semi-automated segmentation strategy in zoom microtomography was able to identify segmented several microstructures (Figure 5A and B) including the glomeruli, Bowman’s capsule and tubules.

**Figure 5.**
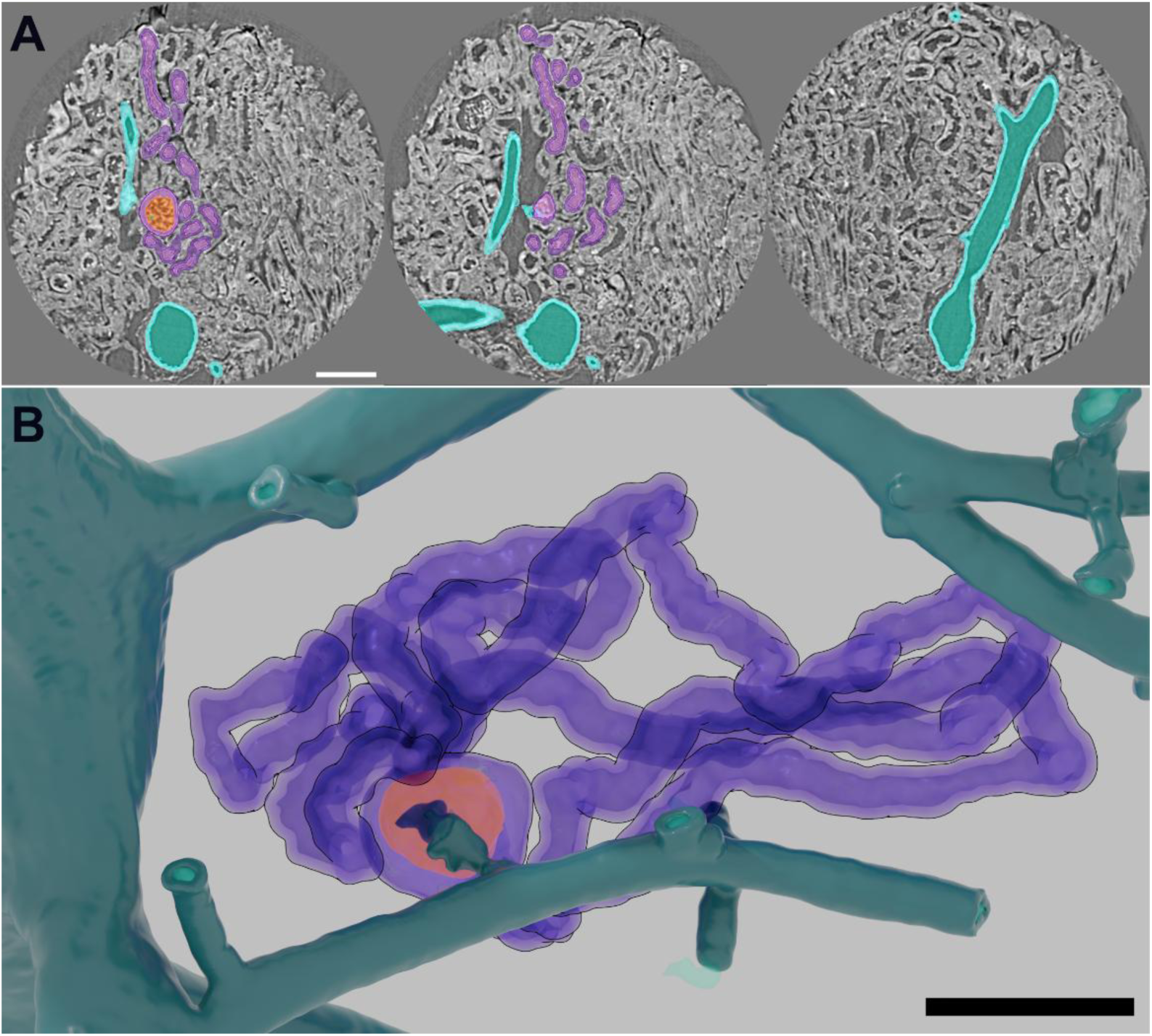
Targeted high-resolution ROI scans from adult murine renal cortex. Panel **A** shows synchrotron 3D tomography slices with segmented structures highlighted: large vessels (light blue), glomerulus (red), Bowman’s capsule and tubules (purple). Panel **B** is volume renderings from segmented structures in same colors as the highlighted virtual slices. Calibration bar = 100 µm.

Furthermore, the non-destructive nature of the micro-CT technique allows the same sample to be used for subsequent conventional 2D histology (Figure 6A and D), enabling direct, point-for-point correlation between the 3D morphological context and immunohistochemical findings. In a small subset of samples (Figure 6B and E), micro-CT scans revealed signs of mild Acute Kidney Injury (AKI^16,17^ with local loss of epithelial integrity, which was not found in other healthy kidneys (Figure 6A and D). AKI findings in synchrotron topographies were further confirmed by conventional 2D histology (Figure 6C and F). AKI histological findings can be associated with chronic kidney disease (CKD) and other conditions that cause decline or loss of renal function^16^. Thus, synchrotron micro-CT overcomes a major limitation of traditional destructive methods by providing details comparable to those of conventional histology within a volumetric context.

**Figure 6.**
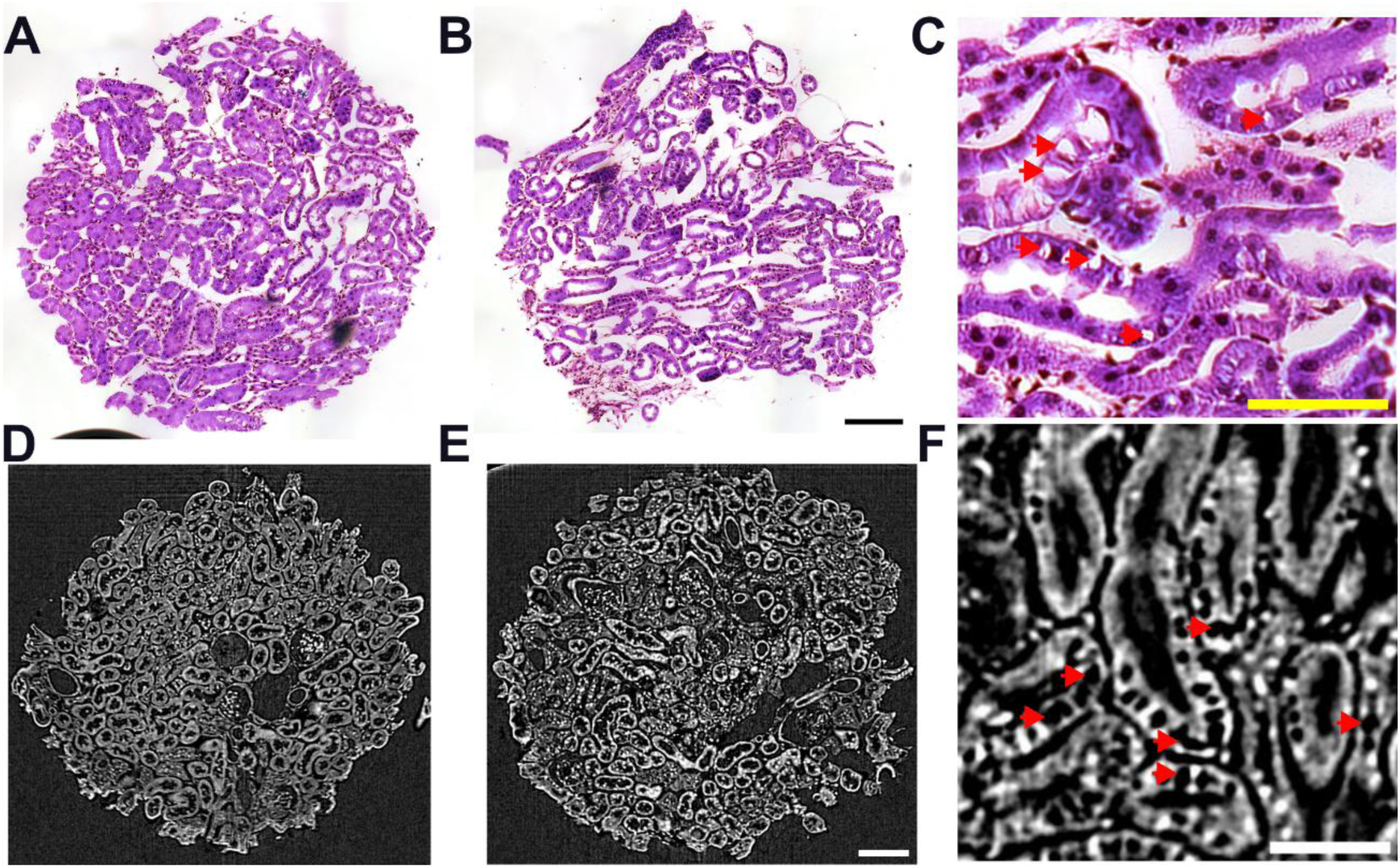
Comparison between synchrotron 3D tomography slices and conventional 2D histology. The images obtained using synchrotron micro-CT has comparable details to conventional 2D histology imaging (**A**,**C**). A small subset of samples (**B**,**E**), micro-CT scans revealed signs of cytoplasmic blebbing in renal tubules, which was further confirmed by conventional 2D histology (**C**,**F**). Calibration bars: 100 µm (**A**,**B**,**D,E**); 50 µm (**C**,**F**).

## 4. DISCUSSION

The successful implementation of this high-resolution micro-CT pipeline provides definitive advantages for renal morphological analysis. Accurate quantification of 3D structures is critical for assessing nephron and microvascular architecture and for providing robust, unbiased measurements for translational studies. Unlike 2D morphometry, which requires large sampling and subsequent stereological extrapolation, the measurements derived from the complete 3D volume eliminates sampling variability. This allows for an objective, comprehensive assessment of the renal microregions that can be spatially heterogeneous. Also, the ability to visualize and quantify this volumetric microstructural structure (with resolution below 500 nm) in relation to the macro-morphology of the organ is unique.

The Mogno beamline at the Sirius Synchrotron proved indispensable for efficiently executing this high-resolution pipeline^11^. Standard laboratory micro-CT devices struggle to achieve the sub-micron resolution needed for histopathology while maintaining a sufficient field of view or minimizing acquisition time^4^. The high X-ray flux available at Mogno enables rapid acquisition (as short as 10 minutes for high-resolution ROI scans), a factor crucial for practical high-throughput preclinical research involving large cohorts and avoiding possible radiation damage to the samples. This efficiency contrasts favorably with benchtop micro-CT machines, which require long scan times are necessary^4,11^.

While the PTA-enhanced absorption contrast method proved effective for visualizing cytoplasmic and interstitial structures and remained compatible with subsequent conventional histology, future technical developments may shift toward X-ray phase-contrast imaging or more targeted contrast agents. Phase-contrast techniques, which are also supported by the Mogno beamline’s design, leverage the refraction of X-rays rather than absorption difference, providing soft-tissue contrast without relying on chemical staining agents^18–21^. Recent work indicates that phase-contrast imaging can yield results comparable to those of contrast-enhanced micro-CT in laboratory settings for kidney imaging^21,22^.

The pipeline developed here provides a standardized, reproducible protocol for non-destructive 3D tissue screening, which is critical for improving the reliability and translational relevance of preclinical nephrology research, especially when assessing drug efficacy or potential toxicity in genetically modified mouse models of CKD^2,8,9,23^. Furthermore, the high volume and unparalleled quality of the 3D data generated by this synchrotron protocol directly address a major hurdle in computational pathology: the lack of high-quality training datasets for 3D biomedical image segmentation^24,25^. Supervised learning algorithms for 3D segmentation often fail to reach performance levels seen in 2D because they lack the necessary annotated volume data for training^26^. By producing more large-scale datasets with validated segmentations of key structures, this pipeline can contribute essential resources for the scientific community to build and expand machine learning investigations, including unsupervised learning, transfer learning, and generative adversarial networks, thereby accelerating the development of generalized AI models for computational pathology by supporting a shift toward standardized, comprehensive 3D data ecology.

This study successfully leveraged the high-flux, multi-scale capabilities of the Mogno beamline at the Sirius Synchrotron to develop a rigorous 3D histology protocol for the murine kidney, utilizing PTA-enhanced X-ray absorption micro-CT. This approach yields objective, quantitative metrics of renal morphology, that overcome the inherent limitations and potential biases of conventional 2D methods. By providing unprecedented spatial and quantitative insight into renal structure, this pipeline serves as a valuable tool for advanced preclinical and translational nephrology research and for high-quality 3D data generation essential for the future of computational histology.

## ACKNOWLEDGEMENTS

This research used facilities of the Brazilian Synchrotron Light Laboratory (LNLS), and the Brazilian Biosciences National Laboratory (LNBio), part of the Brazilian Centre for Research in Energy and Materials (CNPEM), a private non-profit organization under the supervision of the Brazilian Ministry for Science, Technology, and Innovations (MCTI). The Laboratory of Bioimaging (LIB) [proposal 20251190], the Animal Handling and Experimentation Laboratory (LMEA) [proposal 20243527], the Mogno beamline [proposal numbers 20241196 and 20252504] and the Cryogenic Sample Laboratory (LCRIO) facility staff are acknowledged for assistance during the experiments.

---

**Figure V.1.**
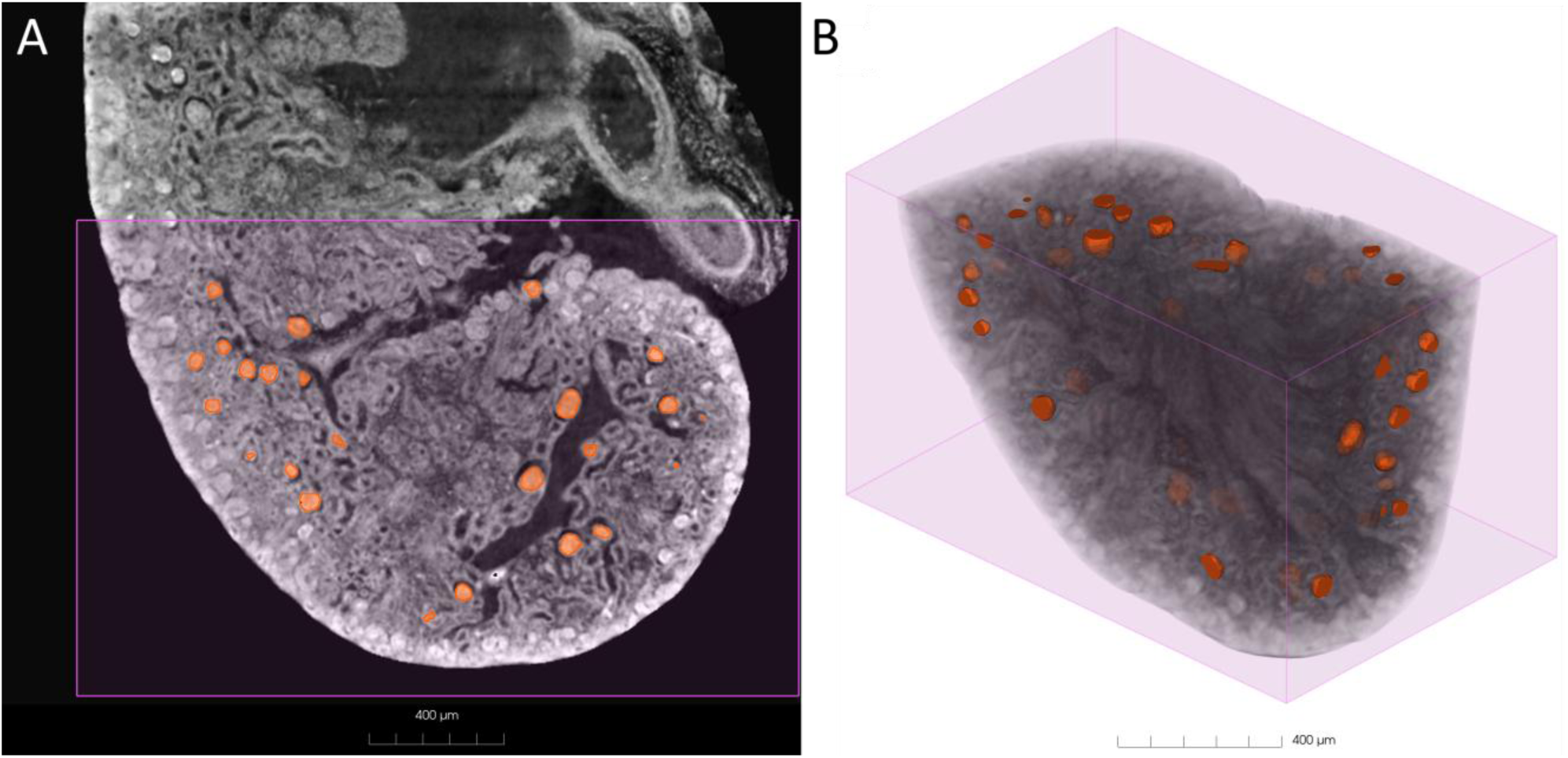
(A) Representative 2D cross-sectional slice of the kidney sample. The purple bounding box indicates the region of interest (ROI) selected for the AI training and inference process. The training labels glomeruli are highlighted in orange. (B) 3D volumetric rendering of the corresponding ROI defined in (A), visualization the spatial distribution of the segmented glomeruli within the tissue structure. Scale bar: 400 µm

**Figure V.2.**
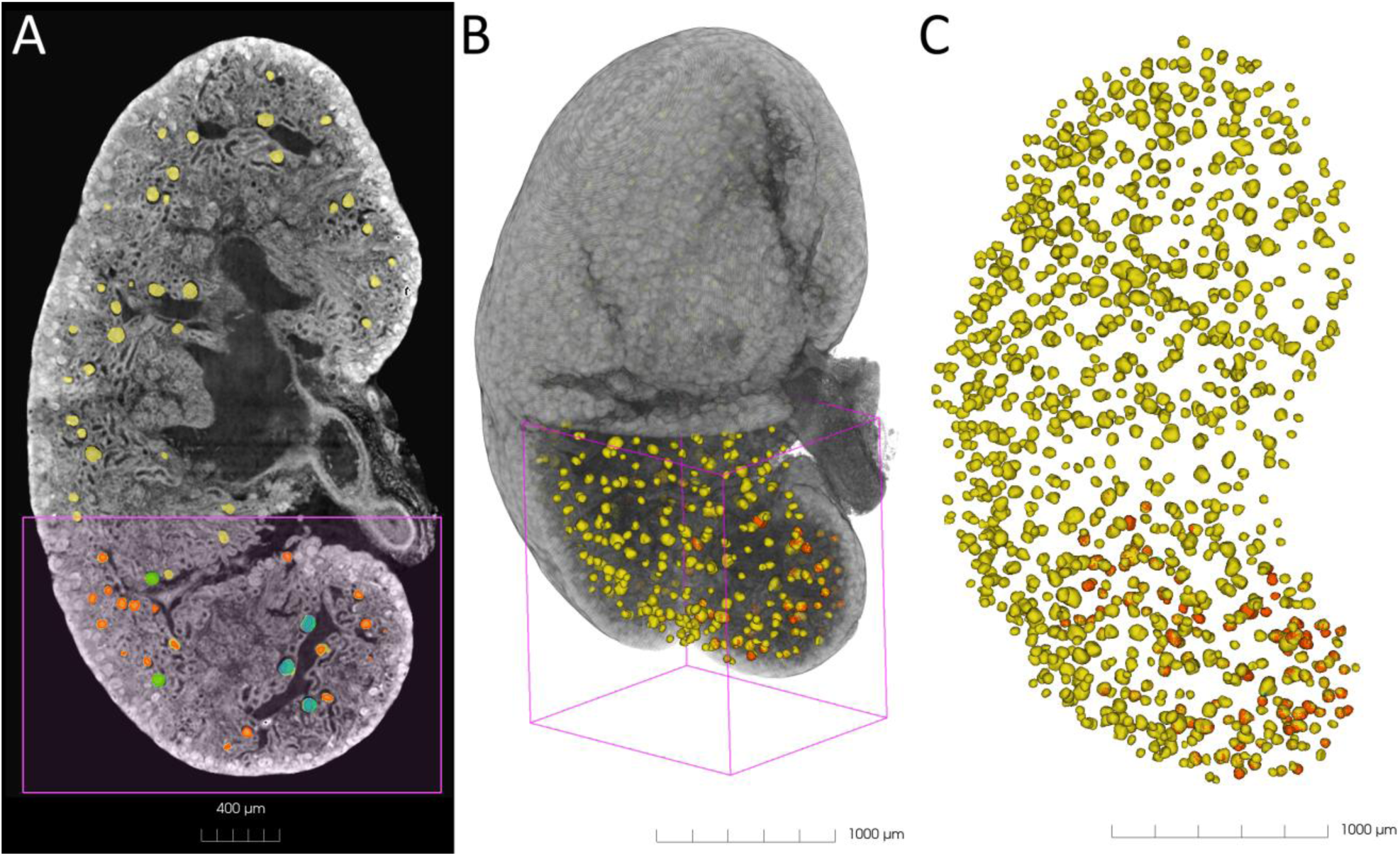
2D cross-section of the entire kidney sample. The purple box highlights the cropped region used for training, while the yellow overlays indicate the glomeruli successfully identified by the AI model in the previously unseen regions (inference). (B) 3D volumetric rendering of the full kidney, demonstrating the model’s ability to generalize the segmentation from the limited training volume (purple wireframe) to the entire organ. (C) Isolated 3D visualization of the segmented glomeruli population, revealing their global spatial distribution and density throughout the sample.

## REFERENCES

1. Busse, M. et al. Three-dimensional virtual histology enabled through cytoplasm-specific X-ray stain for microscopic and nanoscopic computed tomography. Proceedings of the National Academy of Sciences 115, 2293–2298 (2018).

2. Missbach-Guentner, J. et al. 3D virtual histology of murine kidneys –high resolution visualization of pathological alterations by micro computed tomography. Sci Rep 8, 1407 (2018).

3. Metscher, B. D. MicroCT for comparative morphology: simple staining methods allow high-contrast 3D imaging of diverse non-mineralized animal tissues. BMC Physiol 9, 11 (2009).

4. Albers, J., Svetlove, A. & Duke, E. Synchrotron X-ray imaging of soft biological tissues – principles, applications and future prospects. J Cell Sci 137, jcs261953 (2024).

5. Descamps, E. et al. Soft tissue discrimination with contrast agents using micro-CT scanning. Belgian Journal of Zoology 144, (2014).

6. Balcaen, T., Vangrunderbeeck, S., De Borggraeve, W. M. & Kerckhofs, G. Contrast-enhancing staining agents for *ex vivo* contrast-enhanced computed tomography: A review. Tomography of Materials and Structures 7, 100052 (2025).

7. Ougaard, M. K. E. et al. Murine Nephrotoxic Nephritis as a Model of Chronic Kidney Disease. Int J Nephrol 2018, 8424502 (2018).

8. Huang, L., Scarpellini, A., Funck, M., Verderio, E. A. M. & Johnson, T. S. Development of a Chronic Kidney Disease Model in C57BL/6 Mice with Relevance to Human Pathology. Nephron Extra 3, 12–29 (2013).

9. Ehling, J. et al. Quantitative Micro-Computed Tomography Imaging of Vascular Dysfunction in Progressive Kidney Diseases. J Am Soc Nephrol 27, 520–532 (2016).

10. Strotton, M. C. et al. Optimising complementary soft tissue synchrotron X-ray microtomography for reversibly-stained central nervous system samples. Sci Rep 8, 12017 (2018).

11. Archilha, N. L. et al. MOGNO, the nano and microtomography beamline at Sirius, the Brazilian synchrotron light source. J. Phys.: Conf. Ser. 2380, 012123 (2022).

12. Hanly, A. et al. Phosphotungstic acid (PTA) preferentially binds to collagen-rich regions of porcine carotid arteries and human atherosclerotic plaques observed using contrast enhanced micro-computed tomography (CE-µCT). Front. Physiol. 14, (2023).

13. Isensee, F., et al. nnInteractive: Redefining 3D Promptable Segmentation. Preprint at 10.48550/arXiv.2503.08373 (2025).

14. Lösel, P. D. et al. Introducing Biomedisa as an open-source online platform for biomedical image segmentation. Nat Commun 11, 5577 (2020).

15. Lösel, P. & Heuveline, V. Enhancing a diffusion algorithm for 4D image segmentation using local information. in Medical Imaging 2016: Image Processing vol. 9784 707–717 (SPIE, 2016).

16. Gaut, J. P. & Liapis, H. Acute kidney injury pathology and pathophysiology: a retrospective review. Clin Kidney J 14, 526–536 (2021).

17. Nguyen, T. T. U. et al. Deep-learning model for evaluating histopathology of acute renal tubular injury. Sci Rep 14, 9010 (2024).

18. Furlani, M., Riberti, N., Gatto, M. L. & Giuliani, A. High-Resolution Phase-Contrast Tomography on Human Collagenous Tissues: A Comprehensive Review. Tomography 9, 2116–2133 (2023).

19. Momose, A., Takeda, T., Itai, Y. & Hirano, K. Phase–contrast X–ray computed tomography for observing biological soft tissues. Nat Med 2, 473–475 (1996).

20. Quenot, L., Bohic, S. & Brun, E. X-ray Phase Contrast Imaging from Synchrotron to Conventional Sources: A Review of the Existing Techniques for Biological Applications. Applied Sciences 12, (2022).

21. Li, K. Y. C. et al. Feasibility and safety of synchrotron-based X-ray phase contrast imaging as a technique complementary to histopathology analysis. Histochem Cell Biol 160, 377–389 (2023).

22. Mäkinen, H. et al. Comparison of Low-Brilliance X-Ray Phase-Contrast Tomography and Contrast-Enhanced Attenuation-Contrast Micro–Computed Tomography of Rat Kidneys. Kidne*y360* 6, 303–310 (2024).

23. Beeman, S. C. et al. Measuring glomerular number and size in perfused kidneys using MRI. American Journal of Physiology-Renal Physiology 300, F1454–F1457 (2011).

24. Tadokoro, R., Yamada, R. & Kataoka, H. Pre-training Auto-generated Volumetric Shapes for 3D Medical Image Segmentation. in 2023 IEEE/CVF Conference on Computer Vision and Pattern Recognition Workshops (CVPRW) 4740–4745 (2023). doi:10.1109/CVPRW59228.2023.00502.

25. Hu, Y. et al. Artificial intelligence in the task of segmentation and classification of brain metastases images: current challenges and future opportunities. Front. Neurol. 16, (2025).

26. Zhang, L. et al. Generalizing Deep Learning for Medical Image Segmentation to Unseen Domains via Deep Stacked Transformation. IEEE Transactions on Medical Imaging 39, 2531–2540 (2020).

